# Monopolization at the cost of desiccation: Reduced waterproofing cuticular hydrocarbons impairs nestmate discrimination in an ant

**DOI:** 10.1101/2022.10.21.513141

**Authors:** Eyer Pierre-André, Anjel M. Helms, Megan N. Moran, Edward L. Vargo

## Abstract

After humans, social insects represent one of the most complex groups of social organisms, relying on a well-organized communication system among colony members. The transfer of information among individuals is primarily based on cuticular hydrocarbons (CHC). These chemical compounds, produced by all insects, initially evolved to prevent water loss^1^. They were subsequently co-opted as semiochemicals to communicate various types of information. This includes nestmate recognition in social insects^2,3^, enabling different colonies to partition resources by ousting conspecific competitors. In this study, we report the near complete loss of CHC production by workers of the ant *Nylanderia fulva*. This absence of CHCs is a double-edged sword. It represents a causative agent in the ecological success of this ant species — enabling the development of a large supercolony in its invasive range through limited ability to differentiate nestmates— but increases the risk of suffering ecological stress through desiccation.

## Main text

Invasive ant populations are often characterized by a lack of aggression toward non-nestmates, allowing a free exchange of individuals and resources among a network of interconnected nests, a social structure called a supercolony. The supercolonial structure enables invasive populations to reach extreme densities and ecological dominance without intraspecific competition^4^. The development of such a social structure has been suggested to arise from a homogenization of the gestalt CHC odor of distinct nests and, therefore, homogenization of the recognition threshold accepted by workers. This lack of chemical distinctiveness between workers from different nests is thought to originate from the reduced genetic differentiation among nests or the absence of polymorphism at genetic loci influencing chemical profiles^5^. Here, we report the near complete loss of a CHC chemical communication system in a social insect, which may represent a potent catalyst for a route to supercoloniality.

We showed that the quantity of CHC produced by workers of the tawny crazy ant *N. fulva* and its sister species *N. terricola* was drastically reduced compared to CHC levels found in other invasive ants, including smaller workers of the fire ant *Solenopsis invicta* or the supercolonial ant species *Linepithema humile* (Figures 1a, S1; Supplemental information). Although workers of both *Nylanderia* species produced low levels of CHC (X ± SD = 79.0 ± 59.2 ng for *N. terricola* and 74 ± 56.3 ng for *N. fulva*), they exhibit a normal distribution in *N. terricola*, but a bimodal distribution in *N. fulva* (Figure S2). In *N. fulva*, half of the individuals produce a quantity of CHC close to zero (Figure 1c), while the larger quantities found in the other half were mostly, or totally, the result of the production of a single compound (2-Tridecanone; Retention time = 12.82 min; Figures 1d, S3).

**Figure 1:**
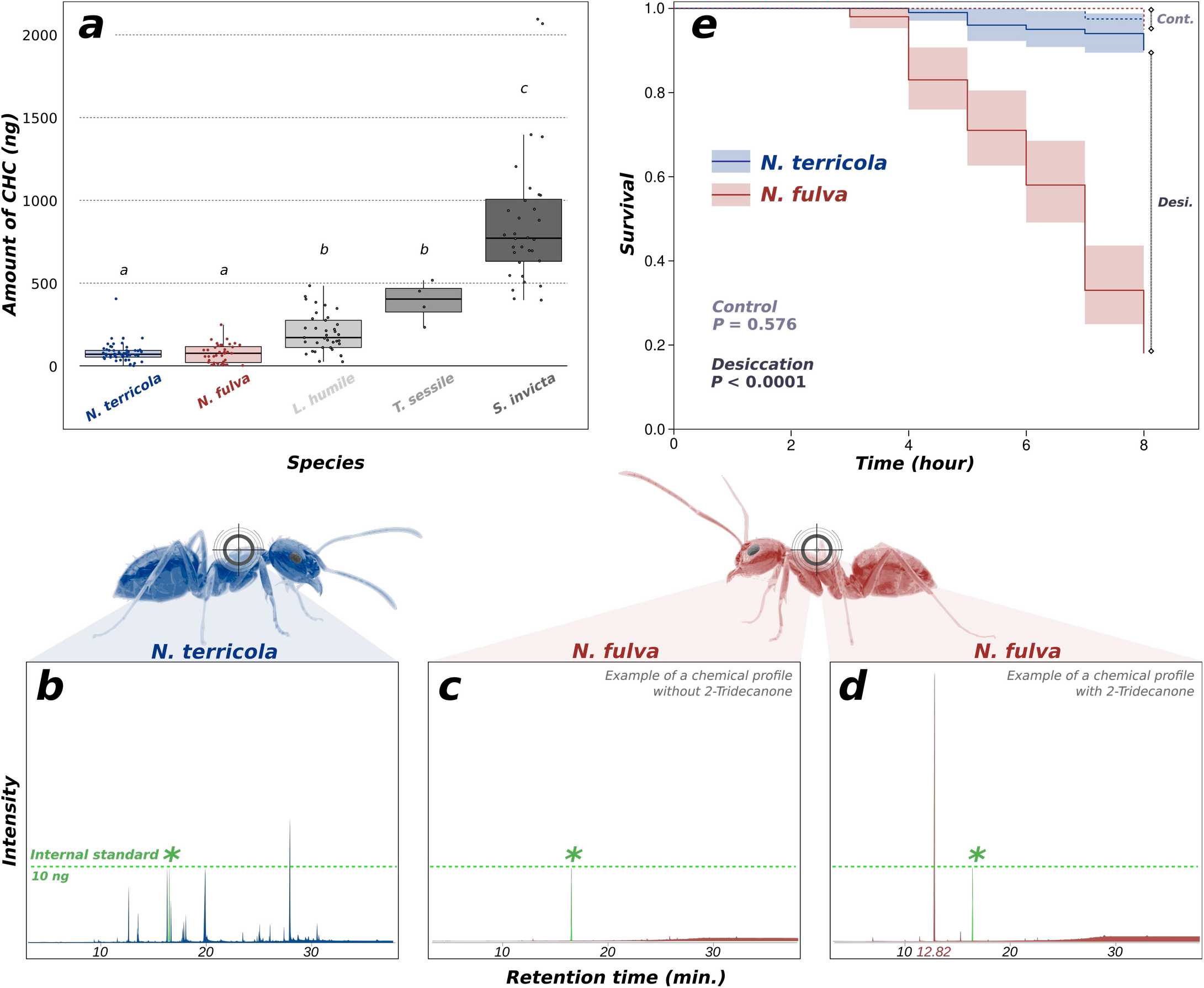
Total amount of cuticular hydrocarbons extracted from individuals of different ant species (*a*). Lowercase letters indicate significant differences among species. Example of cuticular hydrocarbon profiles of workers of *Nylanderia terricola* (*b*), as well as profiles of *N. fulva* without CHC compounds (*c*), or with almost only 2-Tridecanone compounds (*d*). Peaks colored in green and labeled with an asterisk represent the amount of octadecane (C18; internal standard) within samples. Kaplan-Meier survival distributions of workers of *N. fulva* and *N. terricola* for eight hours under desiccation conditions (*i*.*e*., *desi*.; solid lines) and under control conditions (*i*.*e*., *cont*.; dotted lines) (*e*).

2-Tridecanone is a methyl ketone encountered in the Dufour’s glands of many ant species, and abundantly found in those of *N. fulva* (10x more than the closely related species, *Paratrechina longicornis*^6^). Coupled with formic acid, 2-Tridecanone mostly serves as a defensive compound, probably employed by the tawny crazy ant to detoxify *S. invicta* venom during ritualized behaviors^7^. Workers of *N. fulva* grab their acidopore (*i*.*e*., a specialized exocrine-gland duct at the end of the gaster, connected to the Dufour’s gland) with their mandibles and groom themselves vigorously with formic acid and likely 2-Tridecanone. This behavior may explain the large abundance of 2-Tridecanone found on the *N. fulva* cuticle. However, it remains to be determined whether *N. fulva* workers produce other cues not detected by our chemical analyses or secrete additional CHC compounds potentially depleted during frequent acidopore grooming.

As a defensive compound, 2-Tridecanone provides limited information for nestmate recognition (mostly based on methyl-branched alkanes and alkenes^3^).

Therefore, by containing a single, information-poor compound, the chemical profile of *N. fulva* workers does not allow for proper nestmate discrimination. The lack of a strong template to discern non-nestmates has likely favored the development of a supercolonial structure in the introduced range of this species, enhancing its ecological dominance through resource monopolization^8^. This result stands in sharp contrast with other supercolonial invasive ants, such as the Argentine ant, where workers from different nests within a supercolony have similar CHC profiles but there is still sufficient diversity and quantity of compounds to provide a relevant template to identify workers from different supercolonies^9^. The variable aggression levels observed among native nests^10^ may suggest that the information-poor chemical profiles of *N. fulva* has probably acted in concert with a reduction of genetic diversity during its invasion or a gap in the competitive niche of this species in its invasive range to allow the whole invasive population to form a single large supercolony^8^. Evaluating whether the loss of an efficient CHC communication system in *N. fulva* is correlated with changes in the expression of genes associated with CHC production and/or detection, and a relaxed selective pressure on these genes certainly merits further investigation.

Similarly, 2-Tridecanone provides minimal protection against desiccation, which is usually provided by long linear alkanes^3^. We further showed that the reduced production of CHC by *Nylanderia* workers leads to a reduced survival toward desiccation in a low humidity chamber (0-2% RH; Supplemental information) compared to the CHC-abundant species *S. invicta* (Figure S4). All *N. fulva* individuals and ∼70% of *N. terricola* workers died after 16h, while ∼90% of *S. invicta* individuals were still alive at this time. We next demonstrated that the impoverished CHC profiles of *N. fulva* (containing mostly 2-Tridecanone) leads to a reduced resistance toward desiccation. We found that after eight hours in a low humidity chamber, only ∼20% of *N. fulva* individuals survived, while 90% of *N. terricola* workers were still alive (Figure 1e). We confirmed that this difference in mortality is primarily explained by desiccation stress, as 39 out of 40 workers of *N. fulva* (38 out of 40 for *N. terricola*) survived when kept in high humidity chambers for the same time period (control: 100% RH; Figure 1e). The humid and tropical origin of *N. fulva*, and its nesting habitat in the leaf litter have probably lowered the selective pressure toward desiccation stress, therefore relaxing the need for waterproofing efficiency in this species.

Overall, we revealed that the loss of an abundant and diverse CHC mixture has drastic consequences on the ecological success of this invasive ant, affecting both its survival under the stressful environments of the Southern US and its resource monopolization through non-nestmate acceptance. Potentially also a consequence of its capacity to detoxify its major competitor’s venom, we showed that the loss of the CHC communication system and waterproofing system of *N. fulva* represents both a causative agent in its ecological success and its main Achilles’ heel.

## Acknowledgments

We thank the Urban Endowment Fund at Texas A&M University, as well as the Texas A&M AgriLife Research Invasive Ant Research and Management Grant for providing funding for this work.

## Data and code availability

All data generated and analyzed during this study, as well as the code to analyze the data will be deposited upon acceptance in the Open Science Framework database (https://www.osf.io/).

## Supplemental experimental procedures

### Study system

The tawny crazy ant *Nylanderia fulva* is an invasive species, native to South America from Brazil to Argentina, along with Uruguay and Paraguay (Gotzek et al. 2012). This species has first been introduced into Colombia, Peru and the Caribbean (Zenner-Polania 1990; Gotzek et al. 2012; Wetterer & Keularts 2008), but later spread unintentionally to the US. in the 1950’s. Only reported in Florida in the 2000’s, this species has rapidly dispersed to Mississippi, Louisiana, Alabama, Georgia, and Texas in the last two decades (Meyers & Gold 2008; MacGown & Layton 2010; Wang et al. 2016). Genetic data revealed that the invasion of this ant species is associated with founder event(s) significantly reducing the amount of genetic diversity in the invasive range in the US compared to the native range (Eyer et al. 2018). Behavioral and population genetic analyses showed that *N. fulva* displays a multicolonial social structure in its native range, with separate nests maintaining strict boundaries. In contrast, this species exhibits a supercolonial structure in its US invasive range, with no clear boundaries between geographically separated nests (Eyer et al. 2018; LeBrun et al. 2019). Within the southern US, nests are not genetically differentiated from each other. This lack of genetic differentiation is associated with an absence of nestmate recognition. The loss of nestmate discrimination leads to an absence of aggression toward non-nestmates, even from nests separated by hundreds of kilometers, with sharing of individuals and resources between neighboring, interconnected nests (Eyer et al. 2018; Lawson et al. 2020; Kjeldgaard et al. 2020). In addition, each invasive nest is headed by many queens, up to hundreds in number. Overall, these findings show that the entire invasive US range of *N. fulva* is made of a single massive supercolony, extending over more than 2000 km (Eyer et al. 2018, 2019; LeBrun et al. 2019).

### Data collection procedures and chemical analyses

In spring 2021, 40 workers of *N. fulva* were collected in the invasive range of this species in Bryan, TX, US. Individuals were cooled down for one minute on ice before CHCs were extracted by placing each worker into an individual vial containing 200 μL hexane for two minutes. Extracts were evaporated under a stream of high-purity nitrogen and resuspended in 5 μL of hexane including 25 ng of octadecane (C18) used as an internal standard. This solution was transferred to a 100 μL glass conical insert in a 1.5 mL autosampler vial and was analyzed using a 7890B Agilent Gas chromatograph (GC) and 5977B Agilent Mass Spectrometer (MS). A sample volume of 2 μL was injected in splitless mode using a 7693 Agilent autosampler into a HP-5MS UI column (30 m × 0.250 mm internal diameter × 0.25 μm film thickness; Agilent) with ultrahigh-purity helium as the carrier gas (0.75 mL/min constant flow rate). The GC temperature increased from 50 to 310 °C at 15 °C/min after an initial step of 2 min at 50°C and a final held at 310 °C for 10 min. In addition, CHCs were extracted from 52 workers of *N. terricola*, four workers of *Tapinoma sessile*, 32 workers of *Solenopsis invicta*, and 37 workers of *Linepithema humile* following the same procedure. Workers of *Nylanderia fulva, N. terricola, T. sessile* and *L. humile* exhibit a small and continuous size polymorphism, with workers of *N. fulva* being the largest ones (Wild 2004; Gotzek et al. 2012). Workers were randomly chosen for those species. However, *S. invicta* workers exhibit a large variation in body size, with the largest workers being from about 2 to 3-fold as much as the smallest (Tschinkel 2013). Care was taken to only use smaller workers of *S. invicta*, whose sizes are similar to those of *N. fulva* workers (Tschinkel 2013). Overall, the reduction of CHC in *N. fulva* (see Results) unlikely results from an absence of CHC detection, as workers of this species were similar in size or even larger than those of the other species compared (Figure S5).

The abundance of each chemical compound in the CHC profile was inferred from the known amount of internal standard in the sample. The total abundance of CHC was compared between species using the posthoc Dunn test following the Kruskal-Wallis test using the R package *PMCMR*. The P-values were adjusted for multiple comparisons between each pair of species using a Bonferoni correction. For *N. fulva* and *N. terricola*, the distribution of individuals in the overall amount of CHC produced was investigated using the *ggridges* package. All statistical analyses have been performed on R v.3.6.2.

### Desiccation analyses

In spring 2021, nests of *N. fulva, N. terricola* and *S. invicta* were collected in Bryan, TX, US and maintained in the lab for two weeks before the start of desiccation analyses. The colonies were kept under standard rearing conditions (26 ± 2°C, 80 ± 10% relative humidity with LD cycles of 12 h:12 h and fed *ad libitum* with sugar water and cockroaches).

For desiccation analyses, individual ants were placed in Petri dishes with sides coated with Fluon (4.5 cm in diameter). Petri dishes were stacked into covered plastic boxes, above a layer of fully dehydrated Drierite (W.A. Hammond Drierite Co. Ltd., Xenia, OH), decreasing the relative humidity inside the plastic boxes to approximately 0-2%. As a control, Petri dishes with individual ants were placed in plastic boxes without Drierite in 100% humidity chambers. Two experiments were performed. The first one only compared desiccation tolerance between *N. fulva* and *N. terricola*. One hundred individual ants were used from each species under desiccation conditions and 40 individuals from each species were used under control conditions. Worker condition was determined hourly, recording the time to death for each individual. The second experiment compared desiccation tolerance between all three species, 60 individuals were used for *S. invicta* and 24 individuals were used for each of the two *Nylanderia* species. Worker condition was assessed at 16, 24, 40, 48, 64 and 72h, recording the time to death for each individual. In each experiment, the difference in desiccation tolerance between species was visualized using Kaplan-Meier curves and tested using Cox proportional-hazard models (Therneau 2011).

## Supplemental figures

**Figure S1:**
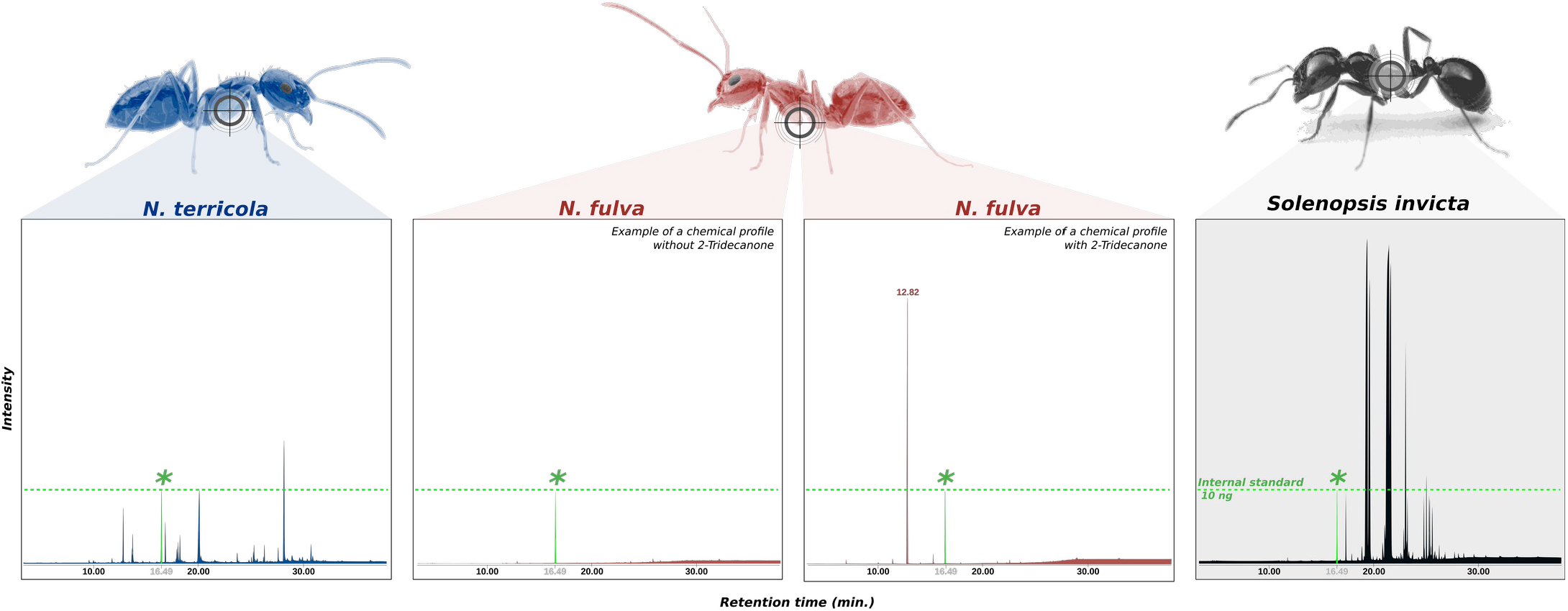
Comparison between cuticular hydrocarbon (CHC) profile of a worker of *Solenopsis invicta* to those of workers of *Nylanderia fulva* and *N. terricola* (also provided in Figure 1b,c,d). Two example profiles are shown for *N. fulva*; each illustrating one example of the bimodal distribution of the levels of CHC observed in this species. One example profile contains almost no CHC compound; the other example profile contains a higher amount of CHC, but almost exclusively composed of the compound 2-Tridecanone. Peaks colored in green and labeled with an asterisk represent the amount of octadecane within samples (C18; internal standard).

**Figure S2:**
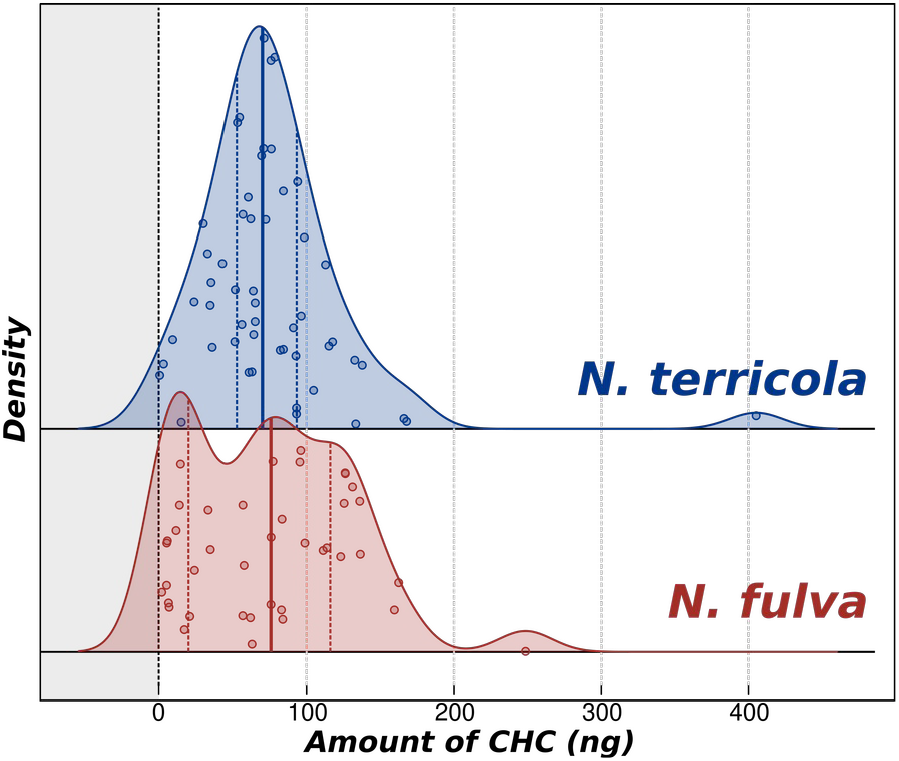
Distribution in the amount of CHCs produced among workers of *N. fulva* and *N. terricola*. Bold solid lines represent the means, dotted lines indicate the 1st and 3rd quartiles for each species.

**Figure S3:**
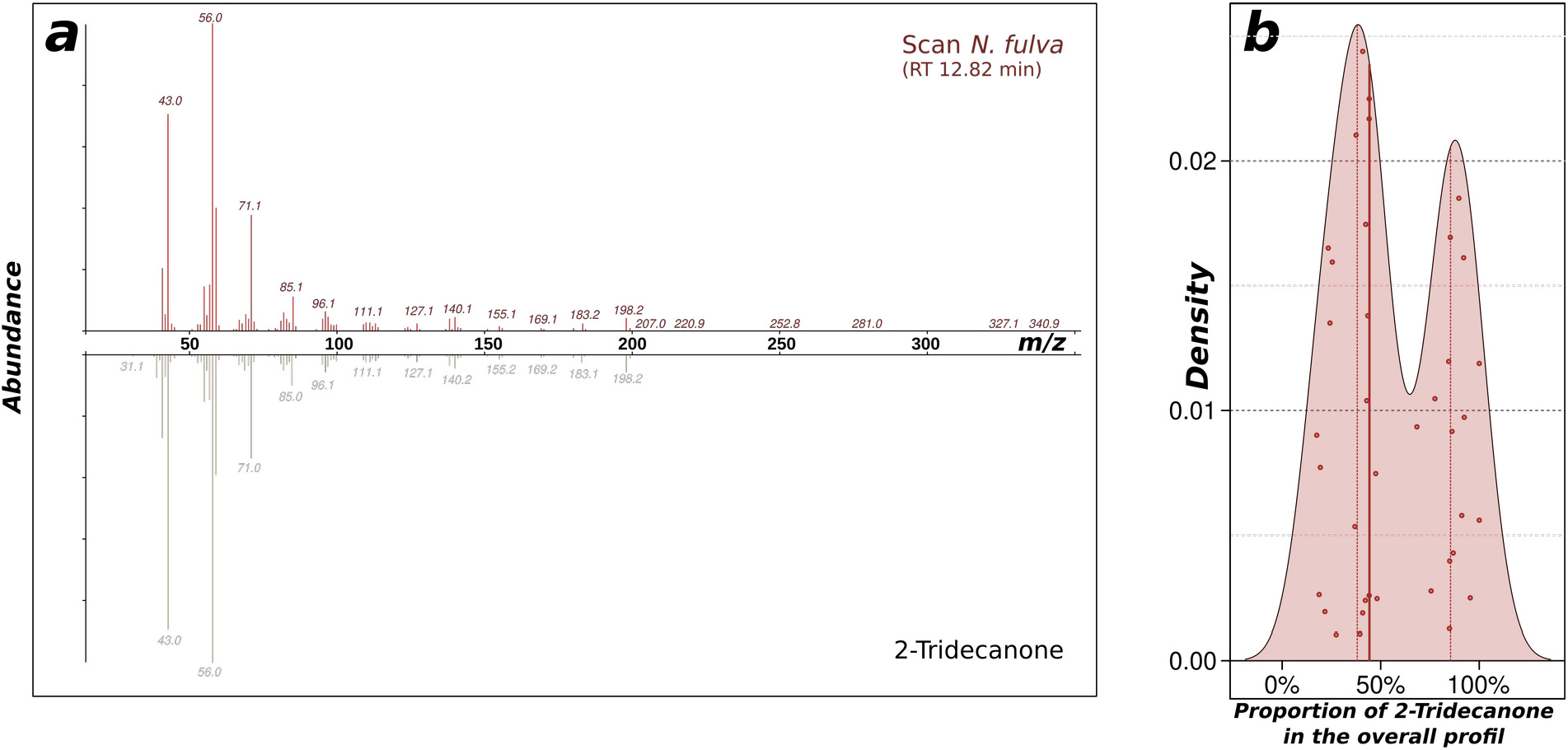
Comparison of the MS fragmentation pathway of the chemical compound found in the *N. fulva* CHC profile at retention time 12.82 min (upper panel) with the expected fragmentation pathway of 2-Tridecanone (lower panel) (*a*). Distribution of the proportion of 2-Tridecanone in the overall CHC profile of *N. fulva* workers (*b*). Bold solid lines represent the means, dotted lines indicate the 1st and 3rd quartiles for each species.

**Figure S4:**
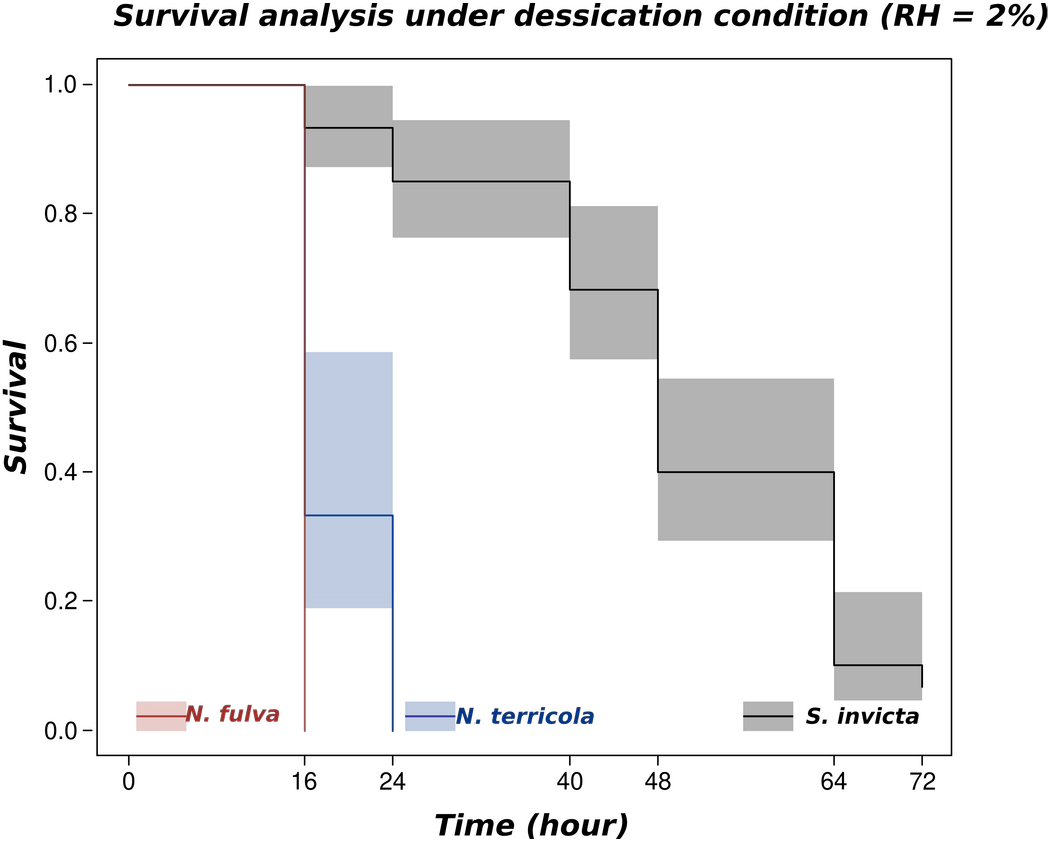
Kaplan-Meier survival distributions of workers of *Nylanderia fulva, N. terricola* and *Solenopsis invicta* for 72 hours under desiccation conditions.

**Figure S5:**
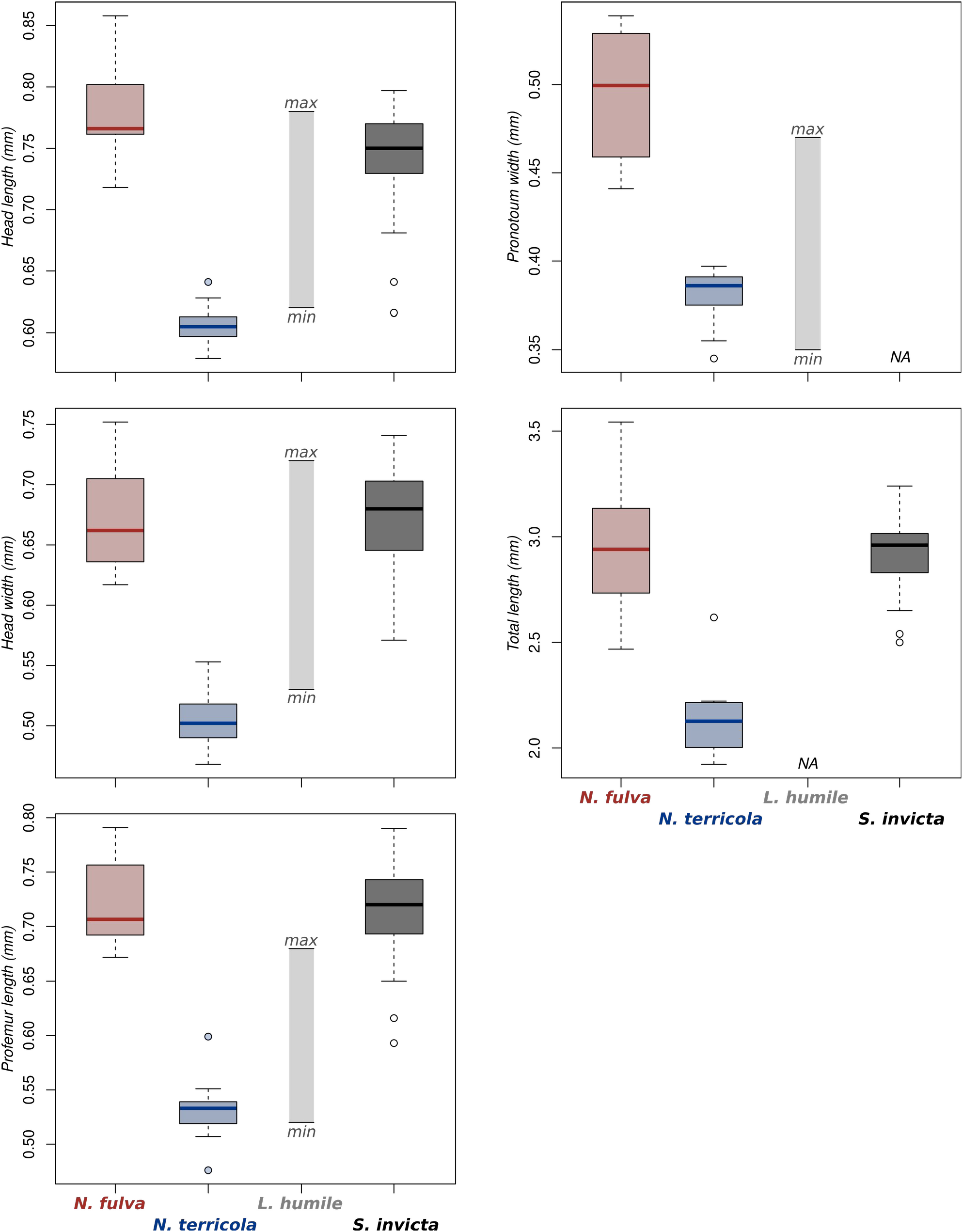
Morphometric comparisons between workers of *Nylanderia fulva, N. terricola, Linepithema humile* and *Solenopsis invicta* for head length and width, profemur length, pronotum width and total length. Morphometric data were extracted from Gotzek et al. (2012, for *N. fulva* and *N. terricola*), Wild (2004, for *L. humile*) and Tschinkel (2013, for *S*.*invicta*). For *S. invicta*, only smaller workers (*i*.*e*., head length < 0.8 mm) were used in this study and were therefore included in the graphs. For *L. humile*, only minimal and maximal values for morphometric measurements were available in Wild (2004). The five different morphometric measurements shown provide a good representation of overall body size in ants (Fournier et al. 2008).

